# A Quantitative Paradigm for Water Assisted Proton Transport Through Proteins and Other Confined Spaces

**DOI:** 10.1101/2021.07.19.452976

**Authors:** Chenghan Li, Gregory A. Voth

## Abstract

Water assisted proton transport through confined spaces influences many phenomena in biomolecular and nanomaterial systems. In such cases, the water molecules that fluctuate in the confined pathways provide the environment and the medium for the hydrated excess proton migration via Grotthuss shuttling. However, a definitive collective variable (CV) that accurately couples the hydration and the connectivity of the proton wire with the proton translocation has remained elusive. To address this important challenge – and thus to define a new quantitative paradigm for facile proton transport in confined spaces – a CV is derived in this work from graph theory, which is verified to accurately describe water wire formation and breakage coupled to the proton translocation in carbon nanotubes and the Cl^−^/H^+^ antiporter protein, ClC-ec1. Significant alterations in the conformations and thermodynamics of water wires are uncovered after introducing an excess proton into them. Large barriers in the proton translocation free energy profiles are found when water wires are defined to be disconnected according to the new CV, even though the pertinent confined space is still reasonably well hydrated and – by the simple measure of the mere existence of a water structure – the proton transport would have been predicted to be facile via that oversimplified measure. In this new paradigm, however, the simple presence of water is not sufficient for inferring proton translocation since an excess proton itself is able to drive hydration and, additionally, the water molecules themselves must be adequately connected to facilitate any successful proton transport.

**Significance Statement:** As first proposed more than 200 years ago by Grotthuss, proton transport is enabled by a chemical bond-breaking and bond-making proton hopping mechanism through water networks or “wires”, often contained within confined systems such as protein channels or nanotubes. Herein, concepts from graph theory are utilized in order to define a new continuously differentiable collective variable (CV) for water wire connectivity and facile proton transport. As such, the water connectivity can be explicitly quantified via free energy sampling, to both qualitatively and quantitatively describe the thermodynamics and kinetics of water-facilitated proton transport via Grotthuss hopping – something that has been lacking since the first conceptual identification of this key chemical process in Nature.

## Introduction

Proton transport (PT) plays a pivotal role in, e.g., the functioning of various biomolecules such as proton exchangers, transporters, and pumps (1–3), as well as certain nanomaterials (4, 5). Stable or at least transient “water wires” are believed to be required for proton permeation through confined regions in these systems by exploiting the Grotthuss proton hopping mechanism (6–8). In Grotthuss hopping the positive charge defect associated with the excess proton in the water structure is transported by dynamically rearranging the chemical-bonding and hydrogen-bonding topology. This process is thus a “chemical” one, involving the breaking and making of chemical bonds. For many years, PT pathways, e.g., in proteins, have been inferred from crystal structures that may show intercalated water molecules and/or from standard molecular dynamics (MD) simulations that study “water wires” in the absence of an explicit proton transport process. Indeed, such MD simulations have been used extensively to provide an atomistic description of water permeation and solvation in nonpolar confined spaces. The presence of water alone has been commonly used (9–17) to predict possible PT pathways, but predicting PT behavior based on hydration alone in the absence of an excess proton in the water structure can be very misleading as is explained below.

It should first be noted that simulating the explicit process of PT through water networks exceeds the capability of traditional MD approaches as widely used in molecular simulations, due to the challenges of modeling the charge delocalization of the excess protonic charge defect and the chemically reactive nature of the excess proton migration (18, 19). *Ab initio* MD, which treats the electronic structure and nuclear motion both explicitly, benefits from its chemically reactive nature, but it can suffer from the shortcomings of insufficiently accurate underlying electronic density functionals and inadequate statistical sampling due to its high computational cost (20). The multiscale reactive molecular dynamics (MS-RMD) method (7, 21–24) (an evolution of the earlier multistate empirical valence bond, or MS-EVB, method) has proven to be capable and successful in simulating PT in a number of biomolecular systems (see, e.g., Refs. (25–31)). MS-RMD (and MS-EVB before it) has a computational efficiency comparable to regular classical MD so that extensive free energy sampling can be carried out for a PT process in large, realistic biomolecular systems.

Indeed, to date relatively few simulation studies have actually included explicit PT behavior, and even fewer experimental measurements have directly probed the PT phenomenon at a detailed molecular level. As such, much speculation has occurred on the nature of proton transport in confined systems such as proteins. Moreover, computer simulations that can treat explicit proton transport have more recently shown a non-trivial coupling of the excess proton translocation to the water hydration of the translocation pathway via a transient and dynamic coupling mechanism (27–35). In essence, it has been found that a hydrated excess proton (hydronium-like) structure can in effect “grow” its own water wire for a subsequent PT process via Grotthuss hopping through that water wire. This occurs because the hydrogen bonding emanating from the excess proton hydronium-like structure is stronger and longer range than normal hydrogen bonding, which can more than compensate for the loss of entropy associated with the formation of the ordered hydration structure (including in a hydrophobic pore or channel). In the majority of these simulation studies, it had appeared to be largely sufficient to simply “count” or “bin” the number of hydrating water molecules in the PT pathway to account for the dynamical coupling of those water structures to the explicit excess proton being present in the water wire (i.e., the water hydration structures are distinctly different – even non-existent – when a hydrated excess proton is not actually in the water wire). A two-dimensional free energy sampling of the excess proton charge migration path in space as one of the coordinates and the degree of hydration (number of water molecules) as the second coordinate has proven especially revealing as to the coupled nature of the charge translocation and water structures in the PT process. Moreover, rate calculations of certain PT processes on this 2D free energy surface has proven to be largely in good agreement with experimental measurements of the proton conduction.

Yet, as can often be the case, a “smoking gun” example emerged (31) that strongly suggested the situation is not so straightforward and that the coupling of PT to hydration requires a more precise and quantitatively powerful description. This particular example involved the ClC-ec1 Cl^−^/H^+^ antiporter protein, but when NO_3_^−^ and SCN^−^ anions are passed through it rather than Cl^−^. The H^+^ transporting activity in these alternative anion cases is significantly disrupted versus the wildtype Cl^−^ limit, and the novel 2.2/1 stoichiometry between chloride and proton transport is also changed. However, the calculation of a 2D free energy surface as described earlier – in this case as a function of excess proton location and the simple hydration of the cavity between the critical E203 and E148 amino acids – provided little explanation for the coupling of the proton antiport to the hydration. Instead, it became clear that the hydration of the PT pathway was simply not enough to account for the overall behavior of the antiporter with an alternative anion flux. And, upon further inspection, it became more obvious that the *connectivity of the water wire hydrogen bonding* – as tied *also* to the location of the excess proton charge defect in the water structure – was a critical feature needed to explain the data. As such, logic suggests that this more complex behavior should be a universal feature of PT processes in confined spaces of proteins and in materials in general, as it certainly includes simple hydration (number of hydrating water molecules) but also goes well beyond just that measure to include structural features (connectivity) of the water wire, and all in the presence of an explicit excess proton.

Though the explicit PT process including the PT-hydration coupling can in principle be simulated in certain cases (27–30, 34) using a method such as MS-RMD (and perhaps direct QM/MM in time), we are still lacking a proper collective variable (CV) to fully describe the hydration component as described above and as directly coupled to the PT, which also often involves the formation and breakage of multiple hydrogen-bonded water networks. Thus, advances in computational methodology in addition to the subtle complexity of the PT processes as described in the paragraph above presents an opportunity to rigorously and quantitatively define a mathematical description of water wire connectivity – as an excess proton is being transported through it – which is continuously differentiable and thus the CV can then be sampled along with the explicit excess proton translocation via an enhanced free energy sampling approach. As such, a proper and general CV can arguably put an end to a great deal of speculation about the relationship between PT processes and internal hydration of confined spaces, e.g., the various speculations and/or partial or incorrect conclusions that have been persisted for many years, largely attributable to missing or incomplete information on the actual PT process.

However, there are several challenges associated with defining and identifying a CV for the purpose stated above. In order to ensure the continuous nature of the CV, a smooth transition in its value is required when waters are entering and leaving the confined space or channel. This issue can be problematic due to the discrete nature of the number of water molecules. Moreover, the CV must be invariant to the water molecule identity and thus be unaffected by frequent exchanges of waters between confined and bulk-like water, and even among the waters within the channel. While systematic CV discovery methods such as the principal component analysis (PCA) (36, 37), time-lagged independent component analysis (tICA) (38, 39), spectral gap optimization of order parameter (SGOOP) (40, 41), auto-encoder/decoder (42–44), Markov state model (MSM) (45), and variational approach for learning Markov processes (VAMP) (46, 47) all represent possible approaches for addressing this problem, each requires an input of possible useful descriptors to be linearly or non-linearly combined into the output CVs. Identifying such descriptors can be more straightforward in some instances; for example, in protein dynamics where the protein conformation can be well described by dihedral angles and contact maps. In contrast, the analogous approach for water network connectivity (especially when it contains an excess proton as is our present focus) is challenging given that there are no *a priori* descriptors available; additionally, the smoothness and identity exchange invariance are also requirements for the input descriptors. More importantly, existing methods typically seek to identify either the largest fluctuating or kinetically slowest degree of freedom, but which provide no guarantee for capturing water network formation or breakage. For example, CVs representing end-to-end distance or the rotation of the entire water wire can potentially be a “learned” output because they are slow motions due to confinement, but they can be minimally (or not at all) related to the PT behavior.

Herein, we present two CVs derived from graph descriptions (48, 49) of water networks, namely the shortest path length (denoted as log(*S*)) and the principal curve connectivity (denoted as *Φ*). We show for contrast that log(*S*), although directly derived from graph theory, is inherently non-differentiable and therefore inappropriate for generating free energy profiles. On the other hand, *Φ*, derived from “coarse-graining” the water graph (cf. Fig. 1A), can serve as a differentiable alternative to log(*S*) and is able to drive efficient free energy sampling of water connectivity via enhanced sampling methods such as umbrella sampling (50) and metadynamics (51). In turn, the principal curve connectivity provides the long-time missing quantitative measure of the transport “capacity” of a given protonated water wire structure for excess protons in a confined space such as a protein channel or a narrow nanotube. Furthermore, the free energy of forming such a structure can now be calculated from this CV and thus the facility of a PT process more readily quantified (i.e., not only speculated upon), as demonstrated below through several examples.

**Fig. 1.**
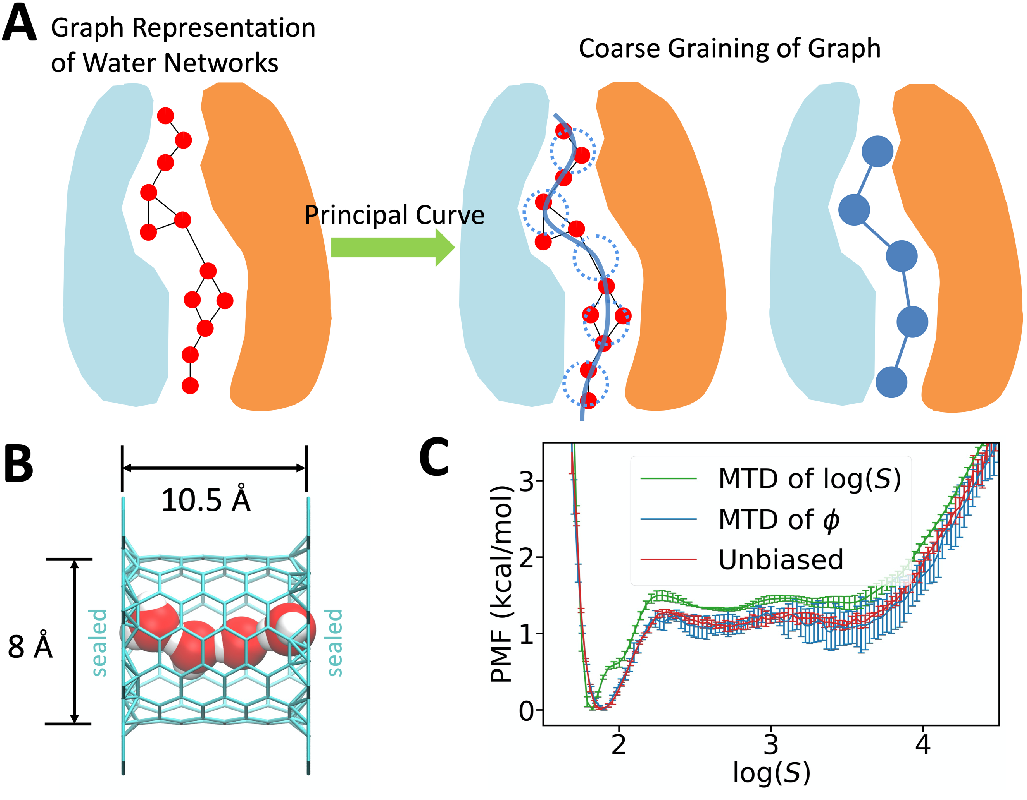
(A) An illustration of water connectivity collective variables (CVs). Water molecules are represented as red spheres. Dotted blue circles along the path represent the “coarse-grained” nodes in the simplified graph. (B) System setup of a sealed carbon nanotube containing 4 SPC/Fw waters. (C) Potential of mean force of the sealed CNT system computed from metadynamics of log(*S*) (green), reweighted from metadynamics of *Φ* (blue), and the reference computed from a long unbiased MD run (red).

In this work, we demonstrate the application of *Φ* to two carbon nanotube (CNT) systems (Fig. 1B and 2B) and a Cl^−^ /H^+^ antiporter, ClC-ec1 (Fig. 3A), from the ClC family (52–56). We combined the MS-RMD method to simulate the explicit proton transport with the *Φ* CV applied to enhance the sampling of the transient water wire connectivity. We show that hydration itself is generally not sufficient for facile PT but, rather, PT primarily occurs when water wires are also fully connected. We further find that excess protons substantially change the water wire conformations and thermodynamics and, consistent with earlier results, that a hydrated excess proton can sometimes create its own water wire where one did not exist before in the absence of the hydrated proton. In doing so the excess proton can reduce the free energy barrier for forming the water wire by ^∼^10 kcal/mol.

**Fig. 2.**
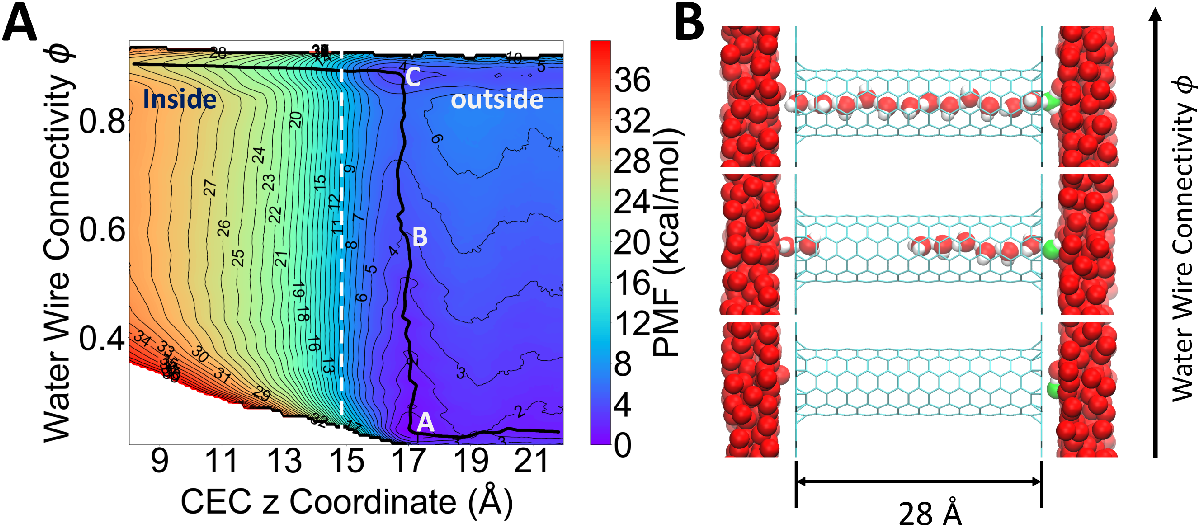
(A) Two-dimensional potential of mean force as a function of *Φ* and CEC z coordinate for proton transport through a hydrophobic nanotube (panel B). The zero point of z-axis is set at the middle of the carbon nanotube and the mouth of the tube is around 14.8 Å in the z position. The minimum free energy path is shown as a black curve. (B) Representative configurations at the positions A, B and C denoted on the 2D-PMF. The most probable hydronium oxygen is shown in green. Hydrogens of water outside the tube are not shown for clarity.

**Fig. 3.**
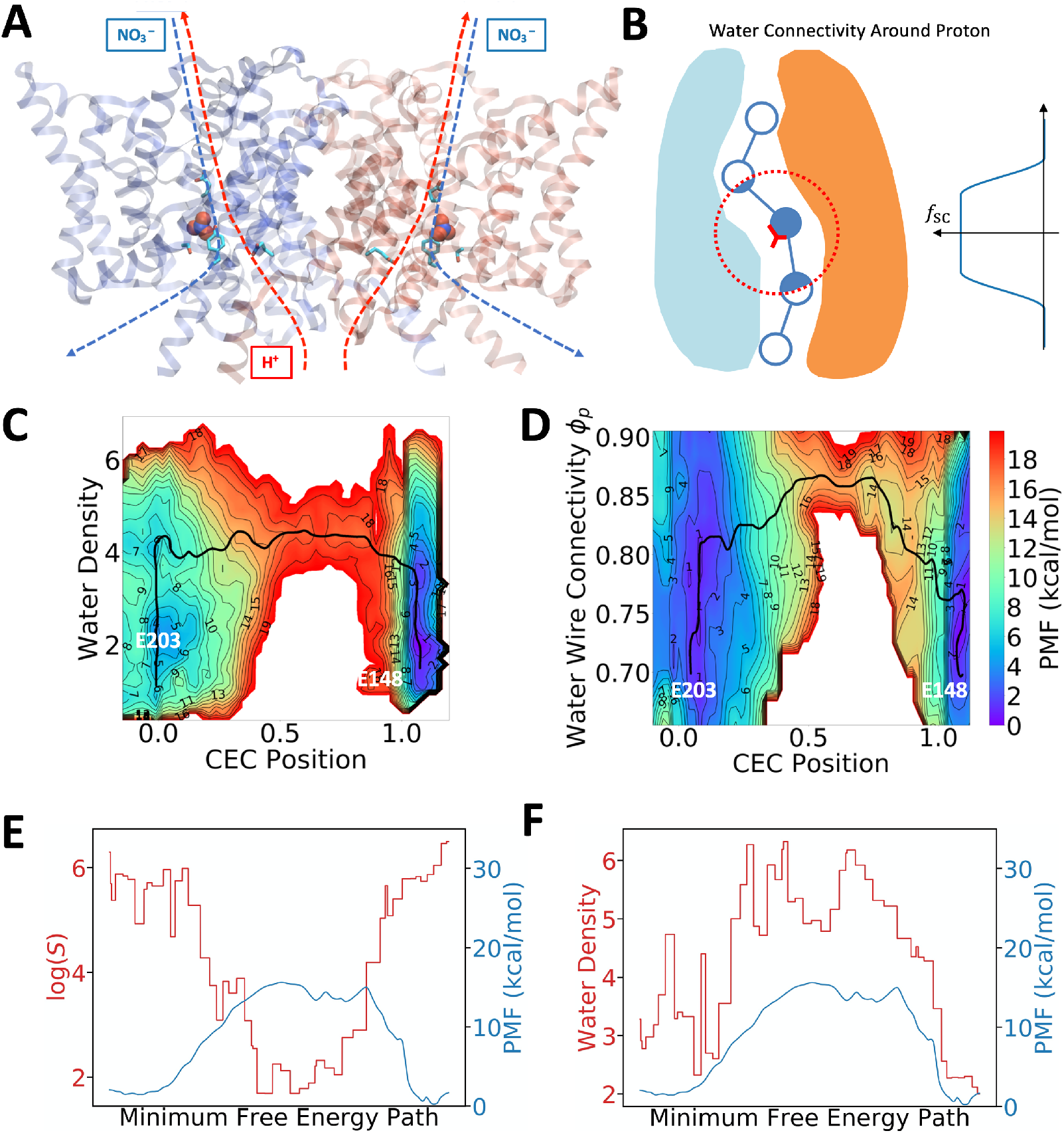
(A) Image of ClC-ec1 in the NO_3_--antiporting state. Proton and anion pathways are indicated by dotted lines. (B) Illustration on how the water connectivity around the excess proton can be calculated. Note that the fictitious beads are spaced more compactly in practice, but the sparse beads here are for clarity of illustration. (C) Two-dimensional potential of mean force (PMF) as a function of simple water density and the hydrated excess proton center of excess charge (CEC) position (Eq. S3) (D) Two-dimensional PMF as a function of water CV, *Φ*, and the hydrated excess proton CEC position (Eq. S3). The black curves shows the minimum free energy path (MFEP) on the 2D PMF (E) The shortest path CV log(S) plotted against PMF along the MFEP (same as black curve in panel B). (F) The water wire gap (see ref (31) for a detailed definition) along the MFEP.

## Methods

### Definition of Shortest Path Length log(*S*)

We consider that each water molecule is a node in a graph (Fig. 1A), and that water networks within a confined system can be fully described by the adjacency matrix of the graph *A*_*ij*_ = *A*_*ji*_ = *f*(*r*_*ij*_), which represents the connectivity between node *i* and node *j* as a function of their distance *r*_*ij*_. The switching function used in this work to approximate infinity when two waters are far apart is

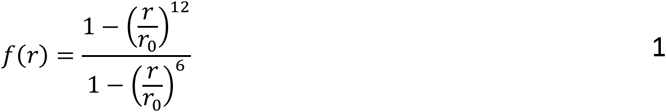

where *r*_0_ = 3 Å as a typical oxygen-oxygen distance of two h-bonded waters. A path on the graph is defined as a collection of connected nodes and thus represents a water chain. We refer to each graph path as a microscopic path because of the atomistic details it provides. We consider that each graph path contributes to the overall connectivity between two given end points via water chains measured by the length of the path (57). The overall connectivity can thus be defined in terms of the shortest path length, indicating the least “effort” required to travel via waters from one end to the other. Since the range of the shortest path length will vary several orders of magnitude due to the nonlinear functions used in *A*_*ij*_, we define the graph connectivity CV as log(*S*), where *S* denotes the shortest path length (Eq. S1 of Supporting Information, SI).

### Definition of Principal Curve Connectivity *Φ*

We consider the water wires to be fluctuating around a 3D curve within the channel (Fig. 1A). By representing an average of microscopic water pathways, this 3D curve describes the “macroscopic” pathway that water is able to permeate. In this way, we effectively “coarse-grain” the atomistic water graph into a simpler graph consisting of a single macroscopic path; accordingly, defining the graph connectivity can be simplified.

We discretize the principal curve into a string of equally separated beads {***x***_*i*_}. A smooth water coordination number for each bead, *s*_*i*_, is calculated to reflect the solvation profile of the curve from a summation of all water oxygens {***x***_*i*_}:

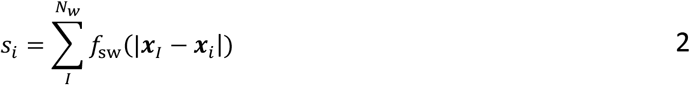

where the switching function *f*_CN_ has the following form

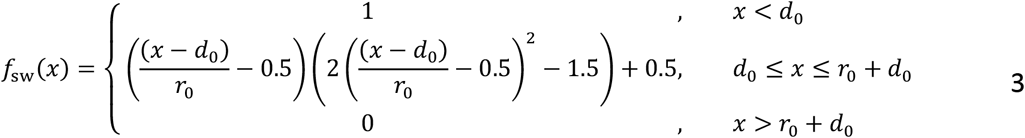

Equation 3 features a plateau when the water deviates from a path bead within *d*_0_ of distance, which allows the water wires to fluctuate around the principal curve but not enough to introduce a change into the final CV value.

Similar to the approach of using an adjacency matrix to describe an atomistic graph, we begin by defining the two-body connectivity in a coarse-grained graph. First, we transform the coordination numbers into the occupancies *I*_*i*_ ranging from 0 to 1 using the Fermi function:

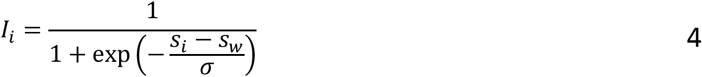

where parameters *s*_*w*_ and *σ* indicate the degree to which a bead along the path is occupied given its water coordination number. Two adjacent beads on the principal curve are connected when they are both occupied, and thus two-body connectivity *f*_*i,i*+1_ can be defined as (*I*_i_ + *I*_*i*+1_)/2. The curve is considered to be fully connected only when all pairs are connected, meaning that the curve connectivity is a logical conjunction (logical AND) of all the two-body connectivity. Based on this fact, we take the product of all *f*_*i,i*+1_ to represent a smooth version of the conjunction to define the final CV as:

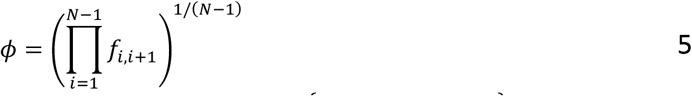

In cases when there are *n* fabricated water pathways in the system, {*Φ*_*j*_ | *j* = 1, 2, …, *n*}, each pathway is represented by a principal curve and is combined by a softmax function to represent the connectivity of the best-connected path:

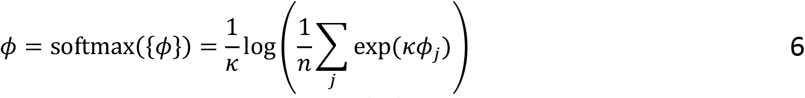

where *κ* > 0 is used to smoothly select the maximum value among {*Φ*_*j*_}. The softmax function here works as a smooth version of logical disjunction (logical OR), meaning that the whole system is allowed to be passed through when any of the pathways are connected. We note that determining an analogous softmin combination of all microscopic path lengths would be computationally formidable because an atomistic water graph contains a factorial number of microscopic paths as a function of the number of waters. Accordingly, we utilize a coarse-grained graph that contains a small number (*n*) of macroscopic paths to overcome this computational impasse.

When the connectivity around the hydrated excess proton is of particular interest, a screening function *a*_*i*_ = *f*_SC_(|***r*** _CEC_ − ***x***_*i*_|) (taking the same form of Eq. 3) can be applied to each bead as the exponent of the occupancy {*I*_*i*_}. The exponent *a*_*i*_ will remain at 1.0 within *d*_0_ of the center of excess charge (CEC) associated with the excess proton charge defect (58), but decays to zero when distant from the CEC, effectively filtering out those less related beads to proton transport (illustrated in Fig. 3B). The resulting connectivity around the hydrated excess proton is denoted as

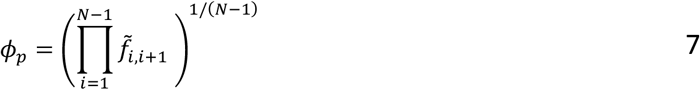

where 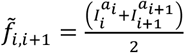.

## Results

### Efficient Free Energy Sampling of Water Wire Connectivity by the Principal Curve Connectivity *Φ*

We first tested our CVs for a model system (Fig. 1B), wherein 4 SPC/Fw waters (59) were sealed in a short carbon nanotube (CNT) by two graphene sheets. The system was designed so that both the connected and disconnected water wires could be effectively sampled by long yet computationally affordable unbiased MD simulations. Hence, these unbiased simulations will provide a reference potential of mean force (PMF) for any CV, specifically the log(*S*) and *Φ* CVs of interest, by directly making histograms. Among the various enhanced free energy sampling methods, metadynamics is known to be a robust tool for studying various chemical and biomolecular processes. The well-tempered metadynamics (WT-MTD) method, which benefits from its asymptotic convergence properties, is one of the most popular variants of the original metadynamics approach (60, 61), and was employed as the enhanced sampling method herein.

Prior to running the metadynamics simulations, we noted from its definition that log(*S*) is not strictly differentiable when two or more paths are the shortest but have the same path length. In other words, log(*S*) becomes non-differentiable when the identity of the shortest path is about to exchange to another path. Since the probability is zero for the system to visit these singularities, one may expect that non-differentiability has only a minimal effect on the free-energy calculation. However, we found a significant deviation of the metadynamics PMF of log(*S*) when compared to the correct one computed from an unbiased molecular dynamics (MD) run (Fig. 1C). We note that the WT-MTD barrier height for connecting water wires for the log(*S*) CV is 1.5 kcal/mol, which is 25% higher than the true (unbiased) value and exceeds statistical error. Additionally, an artificial “shoulder” appears at the position where log(*S*) = 1.8, while only a smooth single well was observed in the unbiased reference. It is therefore striking that biasing log(*S*) resulted in relatively significant errors in such a simple system featuring only 24 permutations for 4 waters. Based on these findings, we eliminated the possibility of using log(*S*) in biased free energy simulations in any real system where the number of permutations is expected to grow factorially as a function of the number of water molecules.

In contrast, a perfectly matching PMF (Fig. S1) was obtained from the WT-MTD of the *Φ* CV which is expected for a differentiable collective variable. Moreover, the correct PMF for log(*S*) was obtained (Fig. 1C) by reweighting the *Φ* metadynamics data using Tiwary and Parrinello’s time-independent estimator (62). This outcome suggests that the new *Φ* CV is a differentiable collective variable appropriate for enhanced free energy sampling, and validates its ability to drive efficient sampling of water wire connectivity.

### Proton Transport and Hydration Coupling in Carbon Nanotubes

The next phase of our investigation involved the use of a more complicated system that was previously utilized to study PT-hydration coupling (32). Specifically, the system consists of two water slabs separated by two layers of inert graphene-like material with a 28-Å-long armchair-type (6,6) single-walled CNT aligned with its z-axis going through the empty space between the two layers. The *ϵ*_LJ_ parameter for carbon was reduced to produce a hydrophobic CNT in the sense that continuous water networks rarely exist in it. To study the proton permeation mechanism through the hydrophobic tube with quantitative accuracy, we carried out two-dimensional umbrella sampling (2D-US) to compute the PMF of (1) the z coordinate of the excess proton CEC and (2) the water connectivity *Φ*. Fig. 2A shows the resulting 2D-PMF and the minimum free energy path (MFEP), revealing a 3-step PT mechanism. First, the hydrated excess proton is transiently trapped at the surface near the CNT mouth, as shown by Point A in Fig. 2A and the bottom panel of Fig. 2B. Then, continuous water wires are spontaneously formed when the excess proton is at the mouth of the tube, as shown by the almost vertical transition A→B→C on the 2D-PMF along the *Φ* direction, and from middle to top panel in Fig. 2B. Finally, the proton permeates through the fully connected water wire (Fig. 2B, top panel) following the horizontal valley on the PMF near the top. Interestingly, this free energy trough can only be seen when the water wire is fully connected corresponding to a *Φ* of ^∼^ 0.9. This is in contrast to any horizontal slice along lower *Φ* values, which correspond to a partially connected wire, and suggests that our newly defined water connectivity plays a critical role for PT through even a simple nano-confined water channel. Moreover, it must be emphasized that the parameters of the CNT were chosen on purpose to make it hydrophobic, so water does not favorably occupy it in the absence of the excess proton; hence, the excess proton “grows” its own water wire for facilitating its transport through the CNT, which is still an energetically uphill (activated rate) process. However, this process has a greatly reduced free energy barrier due to the existence of the transient water wire which facilitates the Grotthuss proton shuttling.

Similar proton-induced hydration was first discovered in previous work (32) by biasing a different hydration CV, namely the simple water occupancy number of the CNT (also referred to earlier as the water density). Though the water density CV, which simply counts the number of waters in CNT, does not directly reflect connectivity as does the present CV, it is not surprising for it to correlate well with connectivity in this simple case because the CNT is modeled as a rigid body that permits only a single file water wire. However, this is not the case for real protein channels and transporters and more complex materials that have other coupled molecular motions and can accommodate the formation of multiple water wires. This is where the connectivity CVs defined in this work will come into play the most, as shown next.

### The Connectivity CV Reveals Proton Transport Coupled Hydration in ClC-ec1

ClC-ec1 is a proton antiporter (Fig. 3A) that also transports anions, such as Br^−^, I^−^, NO_3_^-^, SCN^−^ and Cl^−^, with distinct H^+^ coupling for each anion (63–68). Electrophysiological experiments have determined in descending order the proton coupling for these anions to be: Cl^−^, NO_3_^-^, and SCN^−^. This behavior is indicated by anion-proton stoichiometry ratios of 2.2:1 for Cl^−^ to 7-10:1 for NO_3_^-^, while showing no measurable H^+^ transport for SCN^−^(68). Prior simulation studies (14, 31) have shed some light on the molecular-level activity of the anion modulation mechanism of ClC-ec1 with respect to proton transport. It must be noted, however, that Jiang et al. (14) did not treat proton transport explicitly in their work, and hence did not consider the influence of hydrated excess protons on the transient hydration by water wires. By contrast, Wang et al. (31) did include explicit modeling of excess protons via MS-RMD also with umbrella sampling, but as noted earlier their simple water density CV only provided ancillary evidence for the correlation between PT and water wire connectivity. The results also required an additional qualitative hydrogen bond analysis of the umbrella sampling trajectories in order to better understand the underlying physical behavior. This system is thus the “smoking gun” example alluded to earlier that helped to motivate the search for a quantitative, paradigm-defining analysis of proton transporting water wires presented in this work.

When also sampling the water wire connectivity CV, *Φ*, along with the progress of the hydrated excess proton CEC, the theoretical developments contained herein reveal a significantly complex hydration mechanism arising from coupling with an explicit excess proton. The PT progress from E203 to E148 is revealed from a 2D free energy surface (PMF) calculated in these two CVs, especially as a function of the MFEP on that 2D PMF. Figure 3C shows the 2D PMF of the previous work (31) for ClC-ec1 in its NO_3_ ^−^-antiporting state that biased the simple water density, while Fig. 3D shows the free energy of the same system with the connectivity CV, *Φ*, employed. We first note that in contrast to Fig. 3C which shows more uncorrelated vertical (first) and horizontal (second) motions along the MFEP (black curves in 3C and 3D), the MFEP for the case of using *Φ* features a bell curve-like shape, displaying a simultaneous increase and then decrease in the water wire connectivity *Φ* as the proton transport progresses, as described by the CEC position. This observation clearly shows that the water wire connectivity within the channel is strongly coupled with the excess proton transporting in the water wire (the CEC position in Fig. 3D), which is different from a “PT-occurs-after-water-wire-formation” mechanism that has often been assumed in a great deal of prior literature. The behavior shown here, which can now be clearly demonstrated and quantified by the combination of a reactive MD simulation method involving the actual (explicit) proton transport plus the new water wire connectivity CV, *Φ*, thus calls into question the validity of speculation on PT behavior using only water wire existence alone from crystal structures and/or from standard classical MD trajectories, the latter not including chemical reactivity (i.e., with a fixed bonding topology and hence no possibility of Grotthuss proton shuttling). Likewise, the absence of a pre-existing water wire in a crystal structure or a classical MD simulation *does not necessarily mean that such a wire cannot transiently form*, as a result of a hydrated proton coming into a certain region of a molecular structure and forming its own water wire along the way.

Figure 3E depicts the value of log(*S*) with respect to the MFEP, showing a clear anti-correlation with the 1D PMF extracted along the MFEP. The other two descriptors, namely the probability of forming continuous water wires and the water wire gap length employed in prior work (31), show similar correlations (Fig. S2A and Fig. S2B of the SI). It must be stressed that water wires are fully connected around (and only around) the transition state (smallest value of the path CV log(*S*)), as seen in Figs. 3D, 3E, S2A and S2B, while the simpler water number density already hits high values (~ 4 − 5) roughly at the halfway point of the up-hill part of the MFEP, as shown in Fig. 3F, confirming again that PT not only requires enough hydration of the channel but also requires suitably continuous water wires.

The reaction-rate constants computed from transition state theory (TST) applied to the 2D-PMFs are summarized in Table 1. Note the good agreement with the experimental value (68) relative to the previous simulation in which only the water density CV was used, thus confirming that *Φ* is a better CV for quantifying hydration changes relevant to PT. We also note that in TST, trajectories that cross the reaction barrier are assumed not to cross it again, which is the well-known no-recrossing assumption of TST. In practice, however, using less-than-optimal CVs for the reaction path and dividing surface can omit critical dynamical information about the kinetic rate bottleneck, and TST will result in an overestimation of reaction rates due to the omission of recrossing effects. An optimal set of CVs and dividing surface will minimize the recrossing effects, so in that sense the degree of recrossing may serve as a measure of the quality of a CV when describing an activated rate process of interest. Here, we therefore constructed Markov state kinetic models in the 2D CV space using the dynamic histogram analysis method (DHAM) (69) and computed the reaction rates with recrossing explicitly taken into account by these Markov state models (MSMs) (see SI for details). The MSM reaction rate constants are thus lower than the TST ones, and ratio of MSM to TST defines the transmission coefficient that measures how much recrossing affects the PT reaction. We show in Table 1 that the 2D-US simulation that employed the connectivity CV has a small recrossing effect but the recrossing becomes significant in the case of the simple water density CV, which is just the number of water molecules occupying the channel. For other anion-bound states of ClC-ec1, such as Cl^−^and SCN^−^, we find the recrossing is consistently large when the water density was employed (Table S1). As discussed earlier, our prior study (31) in which water density was employed in a 2D-PMF calculation did not directly reveal a discernible coupling between PT and hydration (Fig. 3C). The dynamics recrossing result again highlights that water density is not an optimal choice for capturing the essential slower motions of water networks as they pertain to proton translocation, which also explains the underestimated reaction rate and low transmission coefficient in that work (Table 1).

**Table 1.**
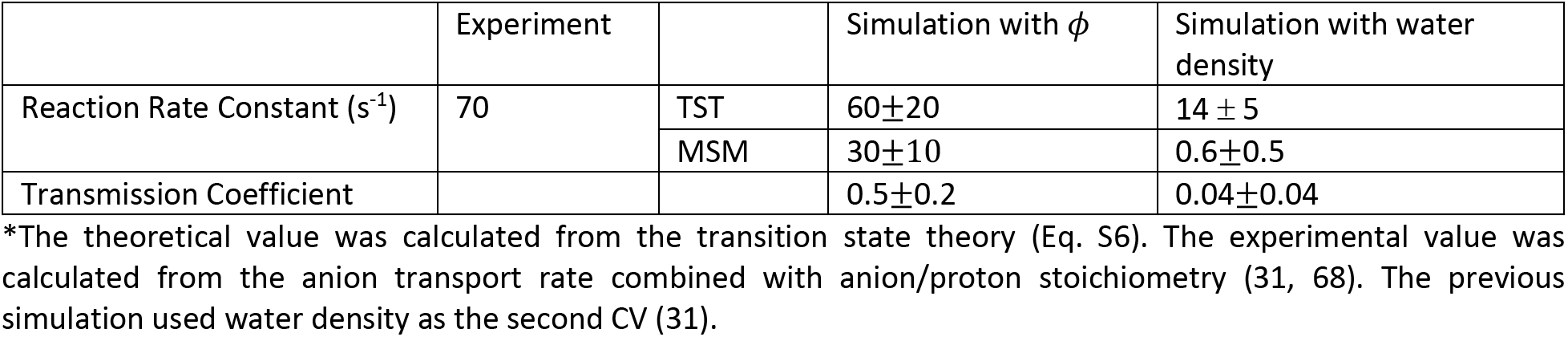
Calculated proton transport rate in ClC-ec1 nitrate-bound state*.

| | Experiment | | Simulation with $\phi$ | Simulation with water density |
| --- | --- | --- | --- | --- |
| Reaction Rate Constant ($\text{s}^{-1}$) | 70 | TST | $60 \pm 20$ | $14 \pm 5$ |
| | | MSM | $30 \pm 10$ | $0.6 \pm 0.5$ |
| Transmission Coefficient | | | $0.5 \pm 0.2$ | $0.04 \pm 0.04$ |

\*The theoretical value was calculated from the transition state theory (Eq. S6). The experimental value was calculated from the anion transport rate combined with anion/proton stoichiometry (31, 68). The previous simulation used water density as the second CV (31).

### Distinct Water Wires in ClC-ec1 Respond to Excess Protons

In order to determine how the hydrated excess proton affects water network connectivity, we performed 1D umbrella sampling of *Φ* with non-reactive classical MD in the absence of an explicit excess proton (Fig. 4A). Fig. S3B of the SI also shows a representative configuration of a single connected water wire obtained from umbrella sampling, consistent with the configurations resulting from unbiased MD (Fig. 4B), thus validating that *Φ* does not introduce artifacts in biased simulations. In contrast to purely classical MD studies, water conformations in the presence of an explicit hydrated proton in reactive MS-RMD simulations show much “broader” water wire structures to facilitate the charge defect delocalization of the excess proton (Fig. 4C). Additionally, reactive simulations with the explicit proton revealed a new water pathway that classical simulations have failed to capture directly, but which was previously suggested by crystal structure analysis (70). These significantly distinct hydration network conformations further highlight the correlation between PT and hydration—namely, that the excess proton not only delivers more solvated water into a hydrophobic channel but can also create new water wires. These findings are also consistent with the observations associated with the CNT model of this work and a prior related study (32).

**Fig. 4.**
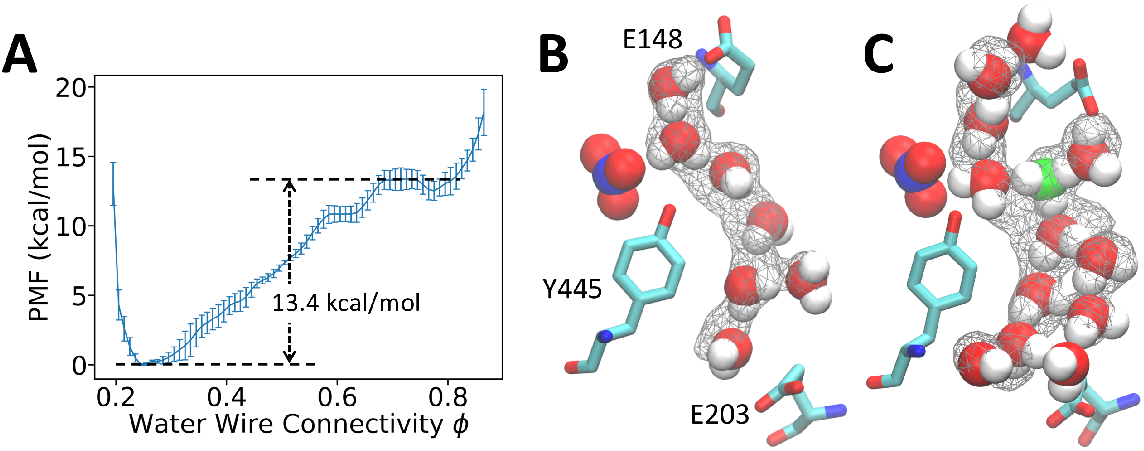
(A) Potential of mean force of *Φ* calculated from standard non-reactive classical MD without an excess proton. (B) A representative configuration of connected water wire sampled in the classical MD. The nitrate anion is rendered in VdW representation. The gray wireframe indicates over 40% water occupancy. (C) A representative configuration of connected water wire sampled with explicitly treating an excess proton by reactive MS-RMD. The most probable hydronium-like structure is shown in green.

The PMF for water connectivity as defined by *Φ* using classical MD is shown in Fig. 4A, which has a water wire connection barrier at 13.4±0.7 kcal/mol. This large barrier makes the formation of fully connected water wires without explicit protons a rare event that requires enhanced sampling to draw accurate quantitative conclusions within affordable computational cost. In order to compare with the cases when an excess proton is free to move (hop or otherwise translocate), we decomposed the total reaction barrier of the 2D-PMF (Fig. 3D) into separate contributions from hydration and from the proton transport according to the gradient theorem,

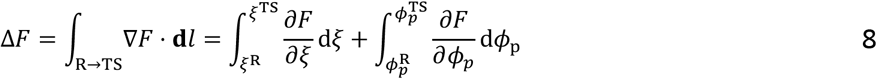

where the integration was evaluated from the reactant minimum to the transition state. The hydration barrier is defined as the integration of mean force in the direction of *Φ*_*p*_:

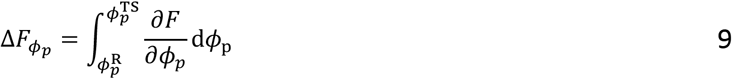

Surprisingly, when an explicit excess proton is present in the water structures, the hydration barrier drops greatly to 1.7 ± 0.8 kcal/mol, which is a few *k*_*B*_^*T*^ at ambient conditions. The origin of the overall free energy barrier for the PT therefore comes more from the electrostatics plus chemical bonding rearrangements of the proton translocation process in the confined region and much less from the cost of forming a water wire alone. When comparing observations with and without an excess proton, the water wire between the E203 and E148 residues was actually mostly disconnected or absent when the excess proton had yet to reach this region; nonetheless, water wires naturally formed as a result of thermal fluctuations during proton transport between the two glutamates. These results again highlight that studies based on standard non-reactive classical MD simulations can be misleading because both the conformations and thermodynamics of the water wires can be substantially impacted by an excess proton being in them.

## Conclusions

In this work we have proposed a new measure (i.e., collective variable, CV) that provides a long sought-after fully quantitative definition of facile water wire connectivity for water-assisted proton transport in confined spaces such as proteins and nanomaterials. We have defined a differentiable CV, *Φ*, that represents water wire connectivity along a principal curve, as well as demonstrated its ability to capture proton transport-coupled hydration dynamics in confined systems. By applying the *Φ* CV to a hydrophobic CNT and a Cl^−^/H^+^ antiporter, and as combined with reactive molecular dynamics and free energy sampling, we are able to identify a novel coupling mechanism between proton transport and water wire connectivity. We find that, even when enough hydration is provided, the hydrated excess proton may often “wait” for fully connected water wires to form before transporting through hydrophobic channels in an activated process fashion. When comparing the reactive MD with classical MD, we also discovered a hydrated excess proton can reduce the free energy barrier of forming continuous water wires by ^∼^10 kcal/mol and can even “create” its own water network. This mathematical description of proton transporting water wire connectivity now provides a powerful and quantitative tool for identifying and characterizing excess proton permeation and hydration coupling behavior in confined spaces such as biomolecular and nanomaterial channels.

## Acknowledgements

The personnel in this research were supported by the National Institute of General Medical Sciences (NIGMS) of the National Institutes of Health (NIH) through grant R01 GM053148. The authors acknowledge the contributions of Dr. Yuxing Peng and Dr. Zhi Wang for kindly sharing their research data and system setup in references (32) and (31). We also thank Dr. Yining Han, Dr. Yuxing Peng, and Dr. Zhi Yue for valuable discussions, and Dr. Paul Calio and Ms. Yu (Grace) Liu for reading the manuscript and providing helpful suggestions. Computational resources were provided by the Extreme Science and Engineering Discovery Environment (XSEDE), which is supported by National Science Foundation Grant OCI-1053575 and the University of Chicago Research Computing Center (RCC).

