## Supplemental File for "A Quantitative Paradigm for Water Assisted Proton Transport Through Proteins and Other Confined Spaces"

**Supporting Figures**

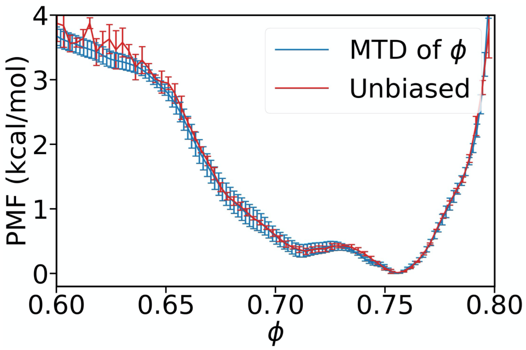


**Fig. S1.** (B) Potential of mean force of $\phi$ in the short CNT computed from well-tempered metadynamics and unbiased molecular dynamics. Note that the small $\log(S)$ value indicates connected water wires, while the large $\phi$ represents connected water wires so that the PMFs of $\log(S)$ and $\phi$ have contrary positions of wells.

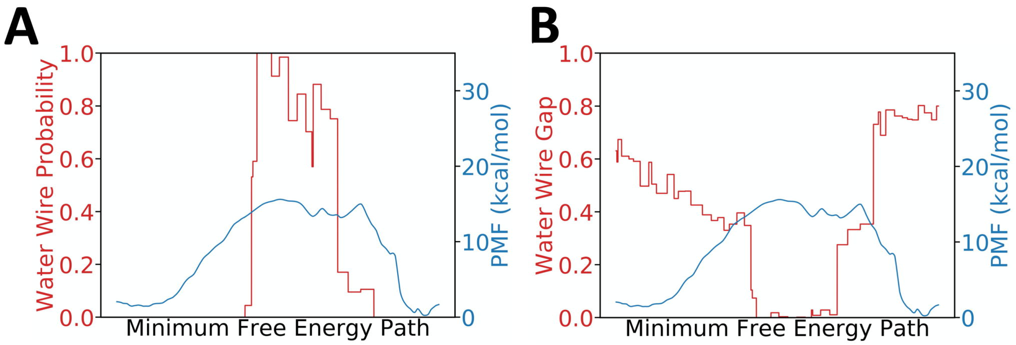


**Fig. S2.** (A) Probability of forming continuous water wire along the minimum free energy path (MFEP). (B) Water density along the MFEP.

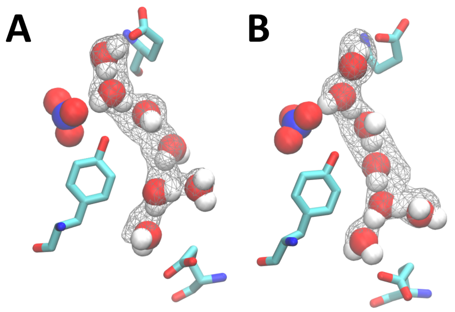


**Fig. S3.** (A) A representative configuration of connected water wire sampled from unbiased MD of ClC-ec1. Note that this figure is identical to Fig. 4B of the main text. (B) A representative configuration of a connected water wire sampled from umbrella sampling of $\phi$. In both figures, the gray wireframe indicates over 40% water occupancy.

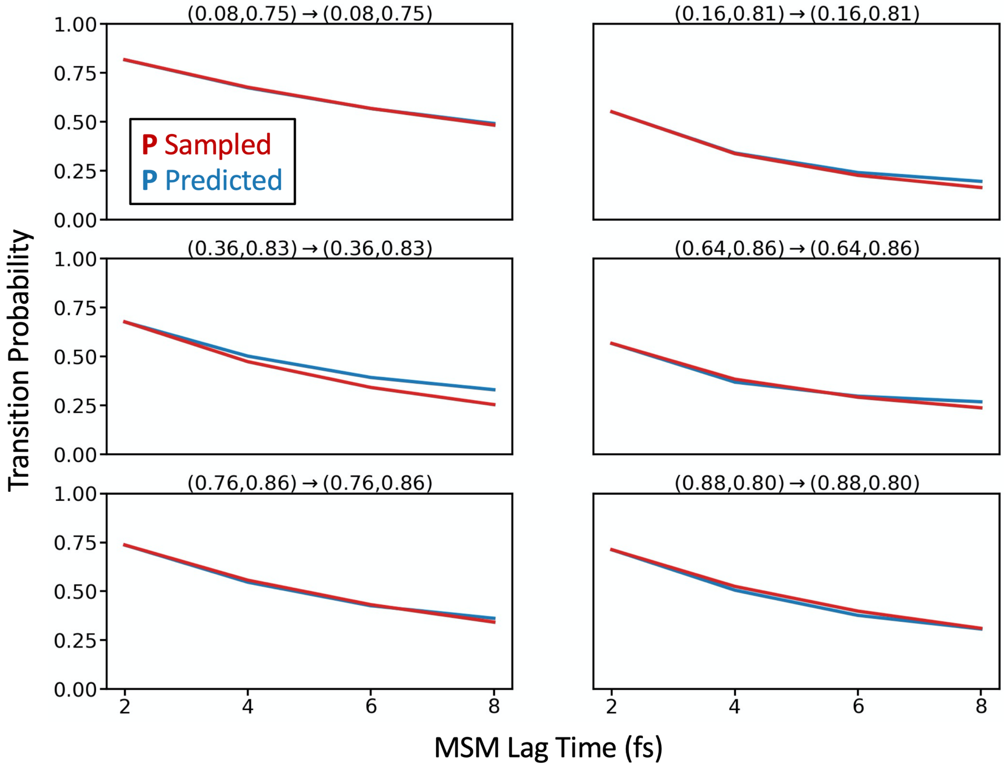


**Fig. S4.** Population of representative microstates along the minimum free energy path. The states are identified as their $(\xi_{\mathrm{CEC}}, \phi_p)$ value.

**Supporting Table**

**Table S1.** Calculated proton transport rates for ClC-ec1 with various bound ions using water density as the hydration reaction coordinate*.

| System | | ClC-ec1 with $Cl^-$ | $SCN^-$ Binding Mode I | $SCN^-$ Binding Mode II |
|---|---|---|---|---|
| Rate Constant ($s^{-1}$) | TST | $(7\pm3)\times10^{7}$ | $(2\pm3)\times10^{-4}$ | $(1\pm2)\times10^{-7}$ |
| | MSM | $(1.27\pm0.01)\times10^{5}$ | $(2.6\pm0.3)\times10^{-7}$ | $(5+4)\times10^{-9}$ |
| Transmission Coefficient | | $(2\pm1)\times10^{-3}$ | $(1\pm2)\times10^{-3}$ | $0.1\pm0.1$ |

*The TST values were taken from ref (1).

### Collective Variable Definitions

Here we provide detailed discussion of the collective variables used in this work, including the choice of their parameters, in addition to the more general descriptions in the main text.

#### Definition of Shortest Path Length $\log(S)$

The adjacency matrix $A_{ij}$ describes the connectivity between any two waters and was given by a switching function in Eq. 1 of the main text. In addition to the specific choice here, we anticipate that any positive-valued, monotonically non-decreasing switching function $f(r)$ that satisfies $\lim_{r\to+\infty} f(r)/r = +\infty$ would work.

Given two nodes $a, b$ on a graph, $S$ is defined as the shortest path length connecting $a$ and $b$:

$$S = \min_{P\in\wp} \sum_{i=1}^{|P|} A_{P_i,P_{i+1}}\ , \tag{S1}$$

where $\wp$ denotes the collection of all paths satisfying $P_1 = a$ and $P_{|P|} = b$. The shortest path is resolved on-the-fly by an implementation (https://github.com/iszczesniak/yen) of Yen's algorithm (2), in the boost graph library (3), and incorporated in PLUMED 2 (4). The end points $a$ and $b$ are chosen to be the carboxyl oxygens of E203 and E148 in ClC-ec1. Due to the absence of protonatable residues in the nanotubes, two virtual atoms located at the center of the circular mouths of the tube are used as end points.

#### Definition of Principal Curve Connectivity $\phi$

In general, the principal curve was computed from the point cloud formed by solvation water using Hastie's algorithm (5). Such was the case in the study of ClC-ec1, but central lines were used for the two CNT systems as they represent the obvious principal curves in rigid straight tubes. The path was discretized at a resolution of 3.2 Å and 2.8 Å in the two CNTs, roughly equivalent to the typical oxygen-oxygen distance of two h-bonded waters. A higher resolution of 1.5 Å was used in ClC-ec1 in order to more accurately capture the curved paths. In the water coordination number (Eqs. 2 and 3), $d_0 = 1.5$ Å and $r_0 = 3.0$ Å for ClC-ec1 were used. A $d_0$ of 3.0 Å was used for the CNTs to address its broader effective width due to the modified Lennard-Jones interaction between carbons and water. The coordination number ranging from 0 to infinity was transformed into the interval $(0,1)$ using the Fermi function (Eq. 4). Parameters $s_w = 1.5$ and $\sigma = 1.0$ were used for the two CNTs, and $s_w = 1.25$ and $\sigma = 2/3$ were used for ClC-ec1. The parameters $d_0 = 5$ Å and $r_0 = 5$ Å for each exponent $a_i = f_{\mathrm{SC}}(\boldsymbol{r}_{\mathrm{CEC}} - \boldsymbol{x}_i)$ were used in ClC-ec1 when defining $\phi_p$ to focus on the water connectivity in near solvation shells of the hydrated excess proton. Generally, it is acceptable to use any parameter for switching functions, as long as the resulting $\phi$ is able to distinguish connected and disconnected water wires. For example, we still obtained the correct $\log(S)$ PMF from reweighting $\phi$ for the short CNT when using the exact same settings as in ClC-ec1, even if the choice of parameters for ClC-ec1 may not be optimal for CNTs. Since a bifurcated path was observed in ClC-ec1, the overall connectivity was combined from the independent connectivity of each of the two paths by a softmax function (Eq. 6) and, to be more specific, was the following for ClC-ec1:

$$\phi = \mathrm{softmax}(\phi_1, \phi_2) = \frac{1}{\kappa}\log\left(\frac{1}{2}(\exp(\kappa\phi_1) + \exp(\kappa\phi_2))\right) \tag{S2}$$

, where $\kappa = 5$ was used to balance smoothness and accuracy of maximum.

#### Reaction Coordinates for Proton Transport

The center of excess charge (CEC) is a virtual "atom" or "site" that tracks the effective position of the net positive charge defect arising from the hydrated excess proton. Its definition and detailed discussion can be found elsewhere (6). The z coordinate of CEC was identified as the obvious coordinate to reflect PT progress along the z-axis-aligned carbon nanotube.

For ClC-ec1, we adopted the same reaction coordinate defined in prior work (1):

$$\xi = \frac{\min_{i\in\{1,2\}} (\boldsymbol{r}_{\mathrm{CEC}} - \boldsymbol{r}_{\mathrm{E203},i}) \cdot \hat{\boldsymbol{n}}_{\mathrm{PT}}}{\min_{i,j\in\{1,2\}} (\boldsymbol{r}_{\mathrm{E148},j} - \boldsymbol{r}_{\mathrm{E203},i}) \cdot \hat{\boldsymbol{n}}_{\mathrm{PT}}} \tag{S3}$$

, where $\hat{\boldsymbol{n}}_{\mathrm{PT}}$ is a predefined unit vector pointing from E203 to E148; $\boldsymbol{r}_{\mathrm{CEC}}$, $\boldsymbol{r}_{\mathrm{E148},i}$ and $\boldsymbol{r}_{\mathrm{E203},j}$ are the positions of CEC, one of the carboxyl oxygens of E148 and one carboxyl oxygen of E203. Using this definition, $\xi = 0$ when E203 is protonated and $\xi = 1$ when E148 is protonated, thus indicating PT progress between the two glutamates. More detailed discussions about this CV can be found in supplemental ref (1).

### Free Energy and Related Calculations

This section provides details regarding enhanced free energy sampling simulations and other related calculations.

#### Metadynamics

The well-tempered metadynamics of $\log(S)$ and $\phi$ both used a Gaussian height 0.025 kcal/mol with a pace of 500 fs. The $\sigma$ of Gaussian was 0.1 for $\log(S)$ and 0.005 for $\phi$ for counting the different scales of the CVs. A bias factor $\gamma = 8$ was used because of a roughly 4 kcal/mol barrier estimated from our unbiased run. The error bars were calculated from block-average by the last 3 blocks of an even partitioning of the full trajectory into 4 blocks.

The PMF of $\log(S)$ under Boltzmann ensemble was recovered from the metadynamics data of $\phi$ by a weighted histogram (7):

$$\mathrm{Prob}(\log S = s^*) = \left\langle \delta(\log S - s^*) \mathrm{e}^{\beta(V(\phi,t) - c(t))} \right\rangle \tag{S4}$$

, where the $\delta$ function was implemented as a Gaussian kernel with a bandwidth of 0.02; the bracket indicates the average under the ensemble generated by metadynamics; $V(\phi, t)$ is the instantaneous bias energy at time $t$ and $c(t)$ was defined as

$$c(t) = \frac{1}{\beta} \log \frac{\int \mathrm{d}\phi \; \mathrm{e}^{-\beta F(\phi)}}{\int \mathrm{d}\phi \; \mathrm{e}^{-\beta(F(\phi) + V(\phi,t))}} \tag{S5}$$

, where the $F(\phi)$ is the PMF of $\phi$ estimated on-the-fly as $F(\phi, t) = -[\gamma/(\gamma - 1)]V(\phi, t)$. Both the metadynamics and reweighting were performed using PLUMED 2.

#### Umbrella Sampling

The umbrella sampling functionality was provided by PLUMED 2 software. The initial configuration of each umbrella window was either the last frame of the umbrella windows in ref (8) and (1), or from equilibration from an adjacent window. In the long CNT, window spacing for CEC z coordinate was 0.5 Å, and as 0.035 for $\phi$, resulting in a total of 420 windows. The force constant for z was 8-10 kcal/mol/Å$^2$ and was 1250-2500 kcal/mol for $\phi$ depending on the curvature of the free energy surface. In ClC-ec1, window spacing for MS-RMD US was 0.025-0.05 for both $\xi$ and $\phi_p$, resulting in 152 windows. The force constant was 1400-4600 kcal/mol and 2500-5000 kcal/mol, respectively. The window spacing of $\phi$ for classical US was 0.025 and the force constant ranged from 2500 to 5000 kcal/mol. All the PMFs were computed from the weighted histogram analysis method (WHAM) (9) using Grossfield's implementation (10). The statistical errors were estimated by block analysis of repeating WHAM on 5 blocks.

#### Rate Constant Calculation

The rate constant was calculated from the 2D-PMF from both the transition state theory (TST) (11), which has demonstrated adequate accuracy in prior studies of ClC-ec1 (12, 13), and a Markov state model (MSM) approach. The TST rate constant is given by

$$k^{\mathrm{TST}}_{\mathrm{E203 \to E148}} = \sqrt{\frac{\langle \dot{\xi}^2 \rangle}{2\pi}} \frac{\int_L \mathrm{e}^{-\beta F(\xi, \phi_p)} \; \mathrm{d}l}{\int_A \mathrm{e}^{-\beta F(\xi, \phi_p)} \; d\xi d\phi_p} \tag{S6}$$

, where $\dot{\xi}$ is the velocity of PT reaction coordinate; $F(\xi, \phi_p)$ denotes the 2D-PMF; the integral in the numerator was evaluated along the dividing curve $L$ at the transition state; and the integral in the denominator was evaluated over the reactant basin $A$ corresponded to E203.

Alternatively, the proton transport rate can be computed from a MSM transition matrix $\mathbf{P} = \{P_{ij}\}$ (14) to account recrossings. A unbiased transition matrix was computed from biased umbrella sampling data by the dynamic histogram analysis method (DHAM) (15), and the Markov property of the resulting MSM was checked by the Chapman-Kolmogorov test (Fig. S4). The forward committor probability $q_i^+$ was first calculated from the transition matrix by solving the following linear equations:

$$q_i^+ = \sum_{i \in I} P_{ik} q_k^+ + \sum_{k \in \mathrm{E148}} P_{ik} \tag{S7}$$

, where $I$ indicates the intermediate region where the excess proton is not bound to either E148 or E203 ($0.3 < \xi_{\mathrm{CEC}} < 0.9$), and the second summation runs over all the microstates that correspond to protonated E418 ($\xi_{\mathrm{CEC}} > 0.9$). For a system in equilibrium, the backward committor probability was computed via $q_i^- = 1 - q_i^+$. The total reaction flux from E203 to E418 was computed from the committors by

$$F = \sum_{i \in \mathrm{E203}} \sum_{j \notin \mathrm{E203}} \pi_i P_{ij} q_j^+ \tag{S8}$$

The reactant state E203 was defined as the region where $\xi_{\mathrm{CEC}} < 0.3$. The proton transport rate constant was computed as the conditional reaction flux per unit time $\tau$ given that the system had visited the E203 state last,

$$k_{\mathrm{E203 \to E148}}^{\mathrm{MSM}} = \frac{F}{\tau \sum_i \pi_i q_i^-} \tag{S9}$$

The MSM rate error was calculated using a bootstrap method in which the trajectories were first partitioned into 2 ps segments, 500 instances of resampling were performed by randomly selecting trajectory segments, and the standard error was computed for the rate constants calculated on the samples. The error in transmission coefficient ($\kappa = k^{\mathrm{MSM}}/k^{\mathrm{TST}}$) was estimated by the error propagation rule

$$\Delta\kappa = \sqrt{\left(\frac{\partial \kappa}{\partial k^{\mathrm{MSM}}} \Delta k^{\mathrm{MSM}}\right)^2 + \left(\frac{\partial \kappa}{\partial k^{\mathrm{TST}}} \Delta k^{\mathrm{TST}}\right)^2} = \frac{1}{k^{\mathrm{TST}^2}} \sqrt{(k^{\mathrm{MSM}} \Delta k^{\mathrm{TST}})^2 + (k^{\mathrm{TST}} \Delta k^{\mathrm{MSM}})^2} \tag{S10}$$

### Simulation Details and System Setup

#### 10-Å-long CNT

The system consisted of 4 SPC/Fw water molecules sealed in a 10-Å-long (6,6) armchair CNT by two layers of 16 Å × 16 Å graphene placed in a 25 Å × 25 Å × 25 Å simulation box. The Lennard-Jones interactions between carbons and hydrogens were set to zero. The LJ parameters between the tube carbons and the water oxygens were $\epsilon = 0.1$ kcal/mol and $\sigma = 3$ Å. The LJ parameters between the graphene carbons and water oxygens were $\epsilon = 0.4$ kcal/mol and $\sigma = 2$ Å. All the carbons were fixed at initial positions and the 4 waters were integrated by a Nose-Hoover chain with a chain length of 3, a timestep of 0.5 fs, and a temperature relaxation time of 250 fs at 310 K. The long-range electrostatic was computed by the particle-particle particle-mesh method (PPPM) with an accuracy of $10^{-4}$. The simulations were carried out with the LAMMPS (16) MD package patched with PLUMED 2. The unbiased run for providing reference free energy profiles was performed for 60 ns. The metadynamics run that biased the shorted path $\log(S)$ was 23 ns. The metadynamics that biased the connectivity $\phi$ was 19 ns and the reweighting was conducted using the last 10-ns of the trajectory.

#### 28-Å-long CNT

The system setup and simulation details were the same as ref (8), except that the MS-EVB 3.2 proton-water model (17) was used instead of the original MS-EVB 3 model (18). The umbrella sampling windows were run for 0.5 ns to 3 ns depending on convergency, resulting in a total of 480 ns of simulation.

#### ClC-ec1

The system setup and MS-RMD simulation details were the same as ref (1). The classical umbrella sampling of $\phi$ was carried out employing the same simulation settings as the MS-RMD simulations, except with the reactive functionality turned off. Each umbrella sampling window time length ranged from 0.3 ns to 4 ns, and the total simulation time was 230 ns.

#### Graphics Details

All the molecular images were rendered by VMD (19). The hydration profile in Fig. 4 and Fig. S3 were computed by VolMap plugin of VMD. All the plots were made by matplotlib (20).
